# Transfer RNA Genes Affect Chromosome Structure and Function via Local Effects

**DOI:** 10.1101/412247

**Authors:** Omar Hamdani, Namrita Dhillon, Tsung-Han S. Hsieh, Takahiro Fujita, Josefina Ocampo, Jacob G. Kirkland, Josh Lawrimore, Tetsuya J. Kobayashi, Brandon Friedman, Derek Fulton, Kenneth Y. Wu, Răzvan V. Chereji, Masaya Oki, Kerry Bloom, David J Clark, Oliver J. Rando, Rohinton T. Kamakaka

## Abstract

The genome is packaged and organized in an ordered, non-random manner and specific chromatin segments contact nuclear substructures to mediate this organization. While transfer RNA genes (tDNAs) are essential for the generation of tRNAs, these loci are also binding sites for transcription factors and architectural proteins and are thought to play an important role in the organization of the genome. In this study, we investigate the role of tDNAs in genomic organization and chromosome function by editing a chromosome so that it lacks any tDNAs. Surprisingly our analyses of this tDNA-less chromosome show that loss of tDNAs does not grossly affect chromosome folding or chromosome tethering. However, loss of tDNAs affects local nucleosome positioning and the binding of SMC proteins at these loci. The absence of tDNAs also leads to changes in centromere clustering and a reduction in the frequency of long range *HML-HMR* heterochromatin clustering. We propose that the tDNAs primarily affect local chromatin structure that result in effects on long-range chromosome architecture.

## Introduction

The three dimensional organization of the yeast nucleus is non-random (Reviewed in (Taddei et al., 2010; Zimmer and Fabre, 2011)). Each chromosome occupies a specific territory in the nucleus anchored to nuclear substructures via specific DNA sequences. The telomeres of each chromosome tend to associate with one another and with the nuclear envelope in small clusters, based on the length of the chromosome arms (Fabre and Spichal, 2014; Palladino et al., 1993b; Ruben et al., 2011). The rDNA repeats on chromosome XII are packaged into a dense structure known as the nucleolus, which also localizes to the nuclear periphery (Oakes et al., 1998). Opposite the nucleolus is the spindle pole body, which is the interphase attachment site for the centromeres of the 16 chromosomes (Jin et al., 2000). Attachment of centromeres to the spindle pole and attachment of telomeres to the nuclear membrane dependent upon chromosome arm length helps organize the nucleus (Therizols et al., 2010). The active genes along the chromosome arms primarily reside in the nuclear interior though some active genes including some tRNA genes interact with nuclear pores and help tether the arms (Duan et al., 2010; Taddei et al., 2010; Tjong et al., 2012).

Besides DNA sequence elements, numerous proteins play a role in nuclear organization via networks of interactions between nuclear membrane and chromatin bound proteins. Chromatin bound proteins involved in this organization include heterochromatin proteins, (Palladino et al., 1993a), lamin like proteins (Andrulis et al., 2002; Bupp et al., 2007; Mekhail et al., 2008; Taddei et al., 2005; Taddei et al., 2004), specific transcription factors (Klein et al., 1992; Taddei et al., 2006), RNA polymerases (Oakes et al., 1998) and DNA repair proteins (Kirkland et al., 2015; Kirkland and Kamakaka, 2013) (see (Taddei et al., 2010) for review).

tRNA genes (tDNAs) are a class of active genes found on all chromosomes and are bound by transcription factors TFIIIB and TFIIIC and RNA polymerase III. tDNAs are short, highly transcribed DNA sequences (Dieci et al., 2007) that are usually nucleosome-free with strongly positioned flanking nucleosomes (Cole et al., 2012; Oki and Kamakaka, 2005; Weiner et al., 2010; Yuan et al., 2005). The tDNAs contain internal promoter elements called A and B-boxes, which aid in the binding of the transcription factor TFIIIC (Geiduschek and Kassavetis, 2001; Schramm and Hernandez, 2002). TFIIIC helps recruit TFIIIB to AT rich sequences upstream of the tDNA. tDNA-bound transcription factors function via interactions with cofactors. tRNA genes are sites of binding for numerous chromatin proteins including the architectural SMC proteins, nuclear pore proteins, chromatin remodelers and histone modifiers. Studies from several labs have shown that tDNAs are enriched in cohesin (Smc1/Smc3) (Glynn et al., 2004), and condensin (Smc2/Smc4) complexes (D’Ambrosio et al., 2008; Haeusler et al., 2008), as well as the SMC loading proteins (Scc2/Scc4) (Kogut et al., 2009; Lopez-Serra et al., 2014) and some chromatin remodelers including RSC (Bausch et al., 2007; D’Ambrosio et al., 2008; Dhillon et al., 2009; Huang and Laurent, 2004; Oki and Kamakaka, 2005).

While individual tRNA genes turn over rapidly as a result of mutational inactivation and gene loss (Frenkel et al., 2004; Goodenbour and Pan, 2006; Withers et al., 2006), a subset of tDNA are syntenic with respect to neighboring sequences (Raab et al., 2012; Wang et al., 2012) and data suggest that these conserved tDNAs possess chromosome position-specific functions in gene regulation (reviewed in (Kirkland et al., 2013; Van Bortle and Corces, 2012)). There are several position-specific effects mediated by tDNAs. First, tDNAs have been shown to function as heterochromatin barrier insulators, which stop the spread of heterochromatic domains into adjacent non-silenced domains (Biswas et al., 2009; Dhillon et al., 2009; Donze et al., 1999; Raab et al., 2012). Second, tDNAs block communication between enhancers and promoters when located between these elements in yeast, *Drosophila*, mouse and human cells by acting as enhancer blockers (Dixon et al., 2012; Ebersole et al., 2011; Korde et al., 2013; Raab et al., 2012; Simms et al., 2008; Simms et al., 2004; Van Bortle et al., 2014). Third, the presence of a tDNA in close proximity to a RNA pol II transcribed gene promoter antagonizes transcription from the pol II transcribed gene in a phenomenon refereed to as tRNA gene mediated silencing (tgm silencing) (Good et al., 2013; Haeusler et al., 2008; Thompson et al., 2003).

In many organisms, tDNAs have also been shown to cluster at sites in the nucleus (Haeusler and Engelke, 2006; Iwasaki et al., 2010; Kirkland et al., 2013; Pombo et al., 1999; Raab et al., 2012). In *S. cerevisiae*, DNA FISH studies have shown that some tDNAs cluster together adjacent to centromeres (Haeusler and Engelke, 2006; Thompson et al., 2003) while proximity ligation analysis suggest that tDNAs cluster at the outer periphery of the nucleolus as well as near the centromeres (Duan et al., 2010) though more recent HiC studies seem unable to detect these long-range associations (Schalbetter bioRxiv 094946). Based on these results it has been proposed that TFIIIC binding to discrete sites along the chromosome plays an important role in chromosome folding and organization in the yeast nucleus (Haeusler and Engelke, 2004, 2006; Iwasaki and Noma, 2012).

To better analyze the role of tDNAs in chromatin looping and organization we generated a “tDNA-less” chromosome through the systematic deletion of all the tDNAs on chromosome III in *S. cerevisiae*. We characterized chromatin packaging, chromosome folding and nuclear dynamics of this chromosome. We show that tDNA loss affects nucleosome positioning and loading of SMC proteins in the vicinity of tDNAs but this has no effect on chromatin looping. While loss of the tDNAs does not affect chromatin looping, it does affect centromere clustering and the long-range interactions of the silenced *HML* and *HMR* loci with concomitant effects on gene silencing.

## Results

The ∼275 tDNAs in the budding yeast genome are dispersed across all 16 chromosomes. Here, we focus on chromosome III, which is 316 kb long and has two tDNAs on the left arm and eight tDNAs on the right arm. In order to investigate the role of tDNAs in chromatin looping, nuclear organization and function, we created a strain in which chromosome III is devoid of any functional tDNAs by deleting an internal fragment of each tDNA. The deletions eliminate the internal promoter elements (both BoxA and BoxB) and thus eliminate the binding of the transcription factors TFIIIC and TFIIIB. For simplicity, we have labeled the tDNA adjacent to the *HMR* locus as t0 and have labeled the remaining nine tDNAs going from right to left as t1, t2, t3 etc. (Figure 1). To delete the tDNAs we first replaced an internal segment of the gene with a *URA3* gene and then subsequently replaced *URA3* with a DNA fragment containing a unique DNA barcode. This involved multiple sequential transformations. Each deletion was monitored by PCR analysis, and intermediate strains were backcrossed to wild type W-303 prior to additional rounds of transformations. All of the experiments described were performed in this strain background to avoid strain specific effects.

**Figure 1.**
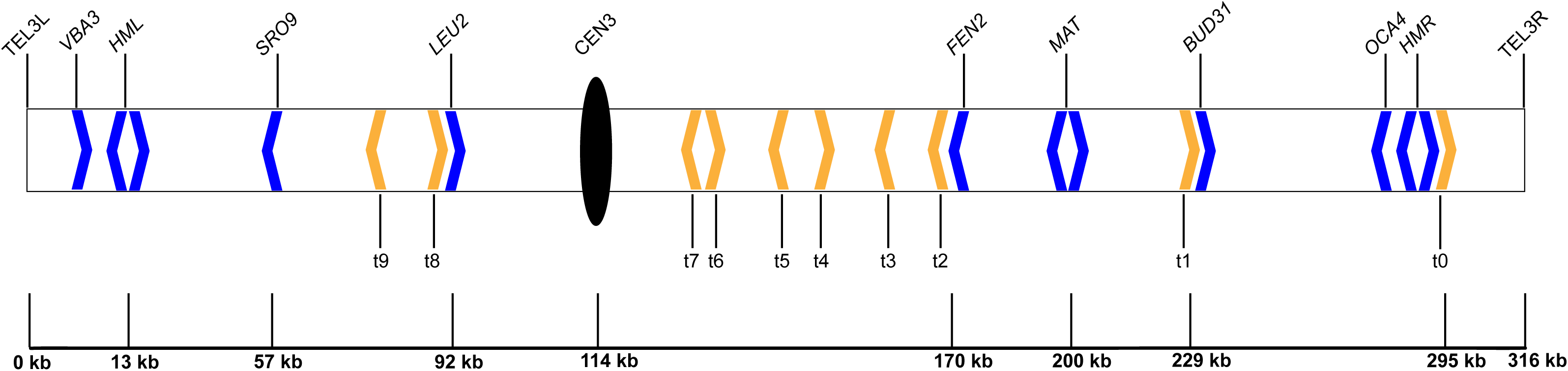
Schematic of Budding Yeast Chromosome III. The positions of the 10 tDNAs on chromosome III are shown. Yellow arrowheads denote tDNAs and the direction of the arrowhead indicates the direction of transcription. Blue arrowheads mark other loci of interest in this study.

**Table 1.** Genes whose mRNA levels changed in the tDNA delete strain compared to the wild type along with statistical analysis of the differences in expression levels.

| **Upregulated in**<br>**tDNA del** | **q-Val (likelihood**<br>**ratio test)** | **q val (Wald**<br>**test)** | **beta**<br>**statistic** |
| --- | --- | --- | --- |
| YCR061w | 0.042003857 | 3.22E-11 | 0.6580998 |
| YDL124w | 0.042003857 | 6.31E-14 | 0.4395261 |
| YHR214c-B | 0.005672182 | 2.35E-210 | 2.5870238 |
| YNL160w | 0.040272092 | 2.69E-20 | 0.5348964 |
| YOR201c | 0.020396347 | 9.18E-27 | 0.7685378 |
| YOR202w | 0.009117145 | 1.20E-76 | 3.7326308 |
| YPL240c | 0.042003857 | 1.20E-15 | 0.5188043 |

| **Downregulated** | **q-Val (likelihood**<br>**ratio test)** | **q val (Wald**<br>**test)** | **beta**<br>**statistic** |
| --- | --- | --- | --- |
| YBR068c | 0.030966243 | 1.53E-20 | -0.5156825 |
| YBR296c | 0.039894621 | 1.87E-19 | -0.5692597 |
| YCR008w | 0.015662463 | 6.48E-43 | -1.1217 |
| YER073w | 0.045197971 | 8.81E-16 | -0.5691673 |
| YER091c | 0.042003857 | 1.79E-20 | -0.5345605 |
| YGL009c | 0.009117145 | 1.47E-82 | -2.0202761 |
| YHR208w | 0.007776891 | 7.72E-106 | -1.3235331 |
| YJR010w | 0.042003857 | 3.34E-13 | -0.8073784 |
| YJR016c | 0.017736878 | 3.09E-29 | -0.5823368 |
| YKL030w | 0.042003857 | 3.46E-11 | -0.6718075 |
| YKL120w | 0.007776891 | 1.47E-94 | -1.4546702 |
| YLR355c | 0.042003857 | 2.35E-12 | -0.387429 |
| YMR108w | 0.042003857 | 3.72E-15 | -0.4213145 |
| YOR271c | 0.027535084 | 4.63E-26 | -0.5903308 |

Most tRNA isoacceptor families have multiple copies, scattered throughout the genome, though single gene copies code for six isoacceptor families. On chromosome III eight of the ten tDNAs that were deleted are members of multi-copy gene families (with 10-16 copies in the genome) and are not essential. However, tDNA t1 (*tS(CGA)c*) is a single copy gene and is essential in *S. cerevisiae* (Ho and Abelson, 1988) and there are only two copies of tDNA t7 (*tP(AGG)c*) in the genome. Loss of t7 from chromosome III caused cells to grow more slowly. In order to remove these two genes from chromosome III and simultaneously maintain the health of the yeast, we integrated single copies of these two genes on chromosome XV at the *HIS3* locus. Once the full tDNA deletion chromosome III had been constructed, the strain harboring this chromosome was backcrossed with wild-type W-303, and segregation of the deleted tDNAs was monitored by PCR using primers specific to the unique barcodes. The full-length sequence of this modified chromosome is available.

The strain where chromosome III lacked any tDNAs (tDNA delete) was grown in rich media at 30C and did not show any obvious growth defect, forming homogeneous and healthy, smooth edged colonies. Strains bearing this tDNA-less chromosome had a doubling time of ∼90 minutes in liquid YPD media, which was indistinguishable from a wild type strain. This is consistent with data showing that loss of one copy of multi-copy tDNAs in yeast cells do not lead to growth defects in rich media (Bloom-Ackermann et al., 2014). Qualitative chromosome loss assays in a homozygous diploid strain, based on the appearance of pseudo-haploids capable of mating, showed no change in chromosome loss rates indicating that chromosome segregation during mitosis had not been perturbed.

### Changes to the local nucleosome landscape surrounding the tDNAs

The stable binding of TFIIIC and TFIIIB as well as their interactions with chromatin remodelers result in nucleosome eviction at the tDNA and positioning of nucleosomes adjacent to the gene (Dion et al., 2007; Oki and Kamakaka, 2005). At some tRNA genes a single nucleosome appears to be disrupted while at other tDNAs multiple nucleosomes are disrupted. Since tDNAs are dispersed across the chromosome and are highly transcribed we first asked if loss of all ten tDNAs from the chromosome altered the nucleosome and transcription landscape of the chromosome. In order to determine if tDNAs affect nucleosome positions across chromosome III, we mapped nucleosomes in our tDNA delete strain as well as in the wild type strain.

Haploid yeast cells were grown to log phase, harvested and nuclei were digested with varying concentrations of micrococcal nuclease to generate mono-nucleosome protected DNAs, which were subjected to paired-end MNase-seq. Overall, the nucleosome landscape across all chromosomes except chromosome III was unaffected by the presence or absence of the chromosome III tDNAs. More focused analysis showed no change in nucleosome positioning in the proximity of the 265 tDNAs scattered on the 15 chromosomes that were not manipulated in this study (Figure 2 left panel).

**Figure 2.**
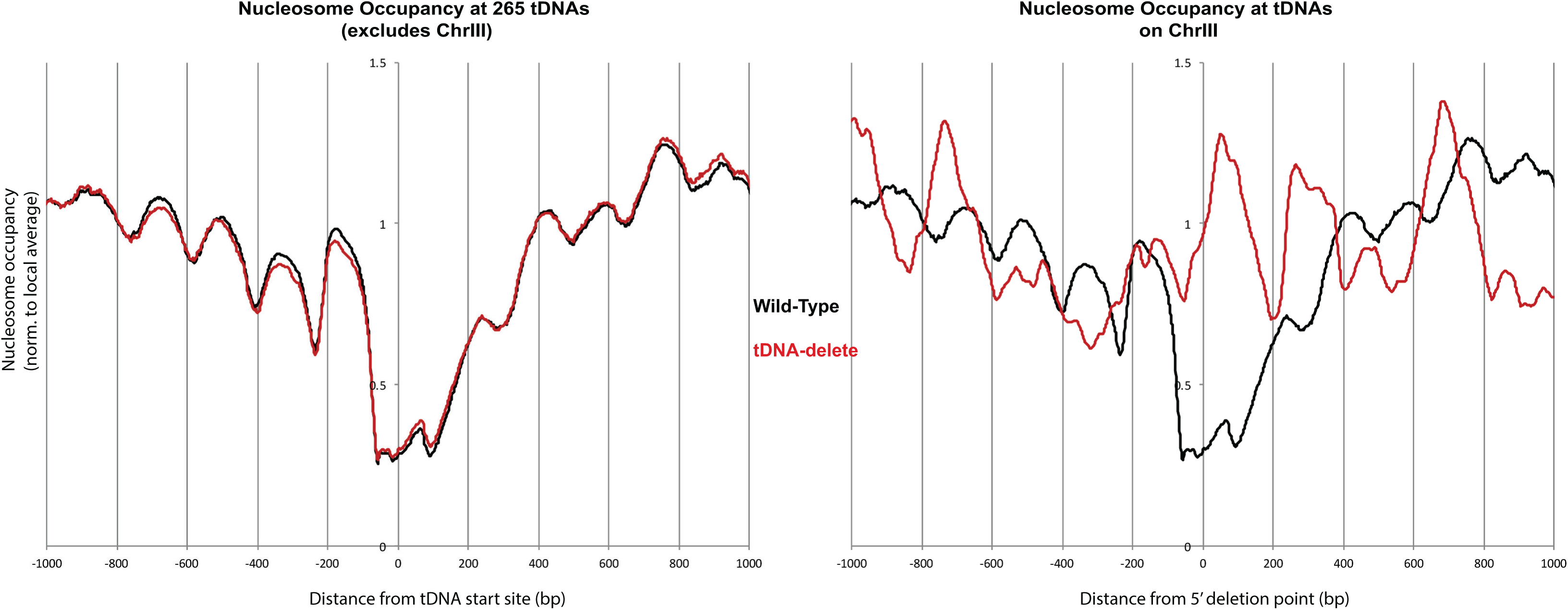
Deletion of tDNAs leads to local changes in chromatin structure. Left Panel: Comparison of nucleosome occupancy at 265 tDNAs on all of yeast chromosomes except chromosome III. The tDNAs were aligned on their transcription start sites (TSS set at 0). WT (black) tDNA delete (red). Right Panel: Analysis of the nucleosome occupancy at tDNAs on chromosome III in the wild type and tDNA delete strain.

In contrast, changes in nucleosome occupancy were observed at or immediately adjacent to the deleted tDNAs on chromosome III. Figure 2 (right panel) shows the average nucleosome occupancy across 2kb segments centered on the chromosome III tDNAs with each tDNA in WT cells aligned at its 5’ end while in the tDNA delete strain, the 5’ ends of the deletion points were aligned. In the wild type strain there is a clear nucleosome free region centered on the tDNA flanked by positioned nucleosomes reflecting differential digestion of the TFIIIB-TFIIIC complex relative to nucleosomes (Chereji et al., 2017). In the tDNA delete strain this pattern is altered and a nucleosome is usually formed over the deletion junction (see Supplementary figure 1). We were unable to determine the change in the chromatin landscape around t1 and t7 tDNAs since these two genes with 100 bp of flanking sequences were transposed to the *HIS3* locus. Nucleosome positions elsewhere on chromosome III that are distant from the tDNAs are not altered on the tDNA-less chromosome (Supplementary figure 2). These results demonstrate that tDNAs create nucleosome free regions at the tRNA gene with positioned nucleosomes flanking the gene. The data also show that their chromatin organizing effects are locally confined and do not extend beyond their immediate vicinity.

### tDNA loss affects expression of very few RNA pol II transcribed genes

The presence of a tDNA in close proximity to a RNA pol II transcribed gene promoter antagonizes transcription from the pol II transcribed gene called tRNA gene mediated silencing (tgm silencing) (Good et al., 2013; Haeusler et al., 2008; Thompson et al., 2003). In addition, tDNAs have also been shown to function as enhancer blockers when located between an UAS enhancer and a promoter (Simms et al., 2008). Since the loss of the tDNAs altered nucleosomes in their vicinity we wondered if these alterations affected the transcription landscape of genes on chromosome III. Rather than restrict the analysis to pol II transcribed genes adjacent to the tDNAs on chromosome III, we investigated the effects of tDNA loss on all pol II transcribed genes in the genome and analyzed the changes in RNA levels in the wild type and tDNA delete strain by RNA-seq. Total RNA was extracted from exponentially growing yeast cultures and RNA-seq libraries were prepared, sequenced and analyzed as described in the materials and methods section. The RNA levels of a very small number of genes were affected upon deletion of the tDNAs. Table1 lists the genes that were either up regulated or down regulated in the strain lacking tDNAs on chromosome III. Of the ten tDNAs present on chromosome III, tDNA t0, t8 and t9 are flanked by retrotransposon elements and since these are repetitive elements, the tDNA-mediated transcription effects could not be investigated for these loci. Furthermore, tDNAs t3 and t4 are missing in W-303. The expression of only two genes on chromosome III was affected and in both instances, a tDNA (t1 and t6) was located adjacent to the gene. In one instance the gene was up regulated upon tDNA loss while in the second instance the gene was down regulated. Furthermore, we observed the up regulation of the *MRM1* gene. This gene resides immediately adjacent to *HIS3*. The tDNAs for t1 and t7 were ectopically inserted at the *HIS3* locus in the tDNA delete strain demonstrating that the ectopic insertion of the tDNAs is the cause of the change in expression of *MRM1*. These data suggest that tDNA mediated position effects are highly context dependent and only affect some pol II transcribed genes and not others.

Of the genes that were down regulated in the tDNA delete strain, several are involved in amino acid biosynthesis though these genes are scattered throughout the genome and do not localize near tDNAs. The reason why expression of these genes was reduced is unclear given that the two yeast strains used are isogenic with respect to nutritional markers, and there are between 10 and 16 copies of each of the six deleted tDNAs in the genome (t0=11 copies, t2=10 copies, t5=16 copies, t6=11 copies, t8=10 copies and t9=15 copies). It is possible that there is a reduction in transcript levels of these genes due to the small reduction in tDNA copy number without any other cell phenotype. This is consistent with a recent study where single tDNAs in yeast were deleted and these single deletions in multi-copy tDNA families also led to changes in the expression of a small set of genes involved in translation (Bloom-Ackermann et al., 2014).

### Scc2 binding at tDNAs is dependent upon a functional tDNA but other binding sites are tDNA independent

The SMC proteins play an important role in nuclear organization (Uhlmann, 2016) and tDNAs are major binding sites for SMC proteins and the SMC loaders Scc2/Scc4 and Rsc. Our nucleosome mapping data indicated that loss of the tDNAs altered nucleosome positions at tDNAs. Since nucleosome free tDNAs are sites for the recruitment of RSC and Scc2/Scc4 proteins (Damelin et al., 2002; Huang and Laurent, 2004; Lopez-Serra et al., 2014; Parnell et al., 2008), we asked if loss of all the tDNAs on chromosome III reduced recruitment of Scc2 proteins at these loci and whether it also affected loading of Scc2 at other sites along the chromosome.

We performed a ChIP-seq of Myc-tagged Scc2 to compare the distribution of this protein genomewide in the WT and tDNA delete strain (Figure 3). This analysis showed that Scc2 bound multiple sites along the chromosome including tDNAs. At some tDNAs the Scc2 binding is focused forming a sharp peak while at other tDNAs the binding is spread over a greater region. Comparison between the wild type and tDNA delete strain showed that Scc2 levels did not decrease at any of the sites on the 15 chromosomes. Upon tDNA loss Scc2 binding decreased at the tDNA loci on chromosome III or at sites in the immediate vicinity of tDNAs such as *LEU2* (adjacent to tDNA t8) (Figure 3 and Supplementary Figure 3) and *HMR* (adjacent to t0). On chromosome III the analysis also showed that there was no significant change in Scc2 binding at other non-tDNA sites. For example we saw a large peak of Scc2 binding at Tel3L. This peak at Tel3L was unchanged upon tDNA deletion (Supplementary Figure4) and similarly we did not record any change in Scc2 levels at *CEN3* (Supplementary Figure4) confirming that tDNAs are not the sole determinants for the recruitment of Scc2 to chromosomes.

**Figure 3.**
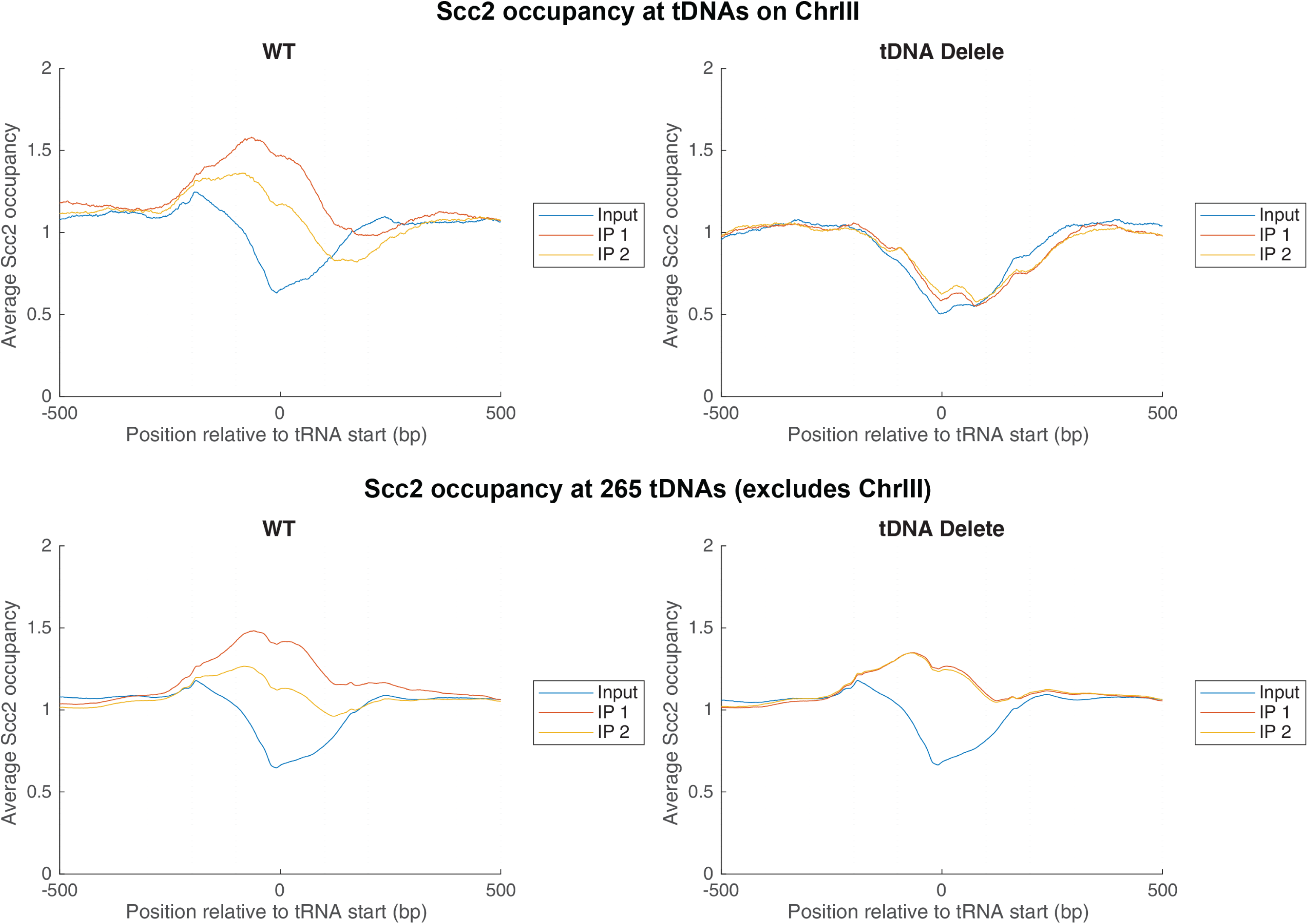
Scc2 binding along chromosome III in the wild type and tDNA delete strain. ChIP-seq mapping of Myc-Scc2. The top panels show the distribution of Scc2 at tDNAs on chromosome III in wild type cells (left) and tDNA delete strains (right). Bottom panels show the distribution of Scc2 at 265 tDNAs on all chromosomes except chromosome III in the wild type (left) and the tDNA delete strain (right).

We confirmed this result by ChIP-qPCR against Scc2. A site at the *OCA4* gene was used as an internal control since this site does not bind Scc2 in wild type cells. We were unable to design unique primers at t6 due to the presence of repetitive sequences in the immediate vicinity of this gene and therefore could not map the localization of these proteins at this tDNA. Some primer pairs flank the tDNAs while others are adjacent to the tDNAs. Consistent with the ChIP-Seq data, in wild type cells, Scc2 is enriched at several of the tDNAs present on chromosome III (Figure 4A). We observed ∼3.5 fold enrichment at t8 and ∼2.5 enrichment at t0, t2 and t5. When the same protein was mapped in the tDNA delete strain we observed a significant reduction in Scc2 binding at these tDNAs. The levels dropped to those observed for the negative control *OCA4* except for the t8 tDNA, where the level dropped two fold but there was some residual Scc2 still present (Figure 4A). The amount of Scc2 did not change at *CEN3* when the tDNAs were absent from the chromosome, indicating that the binding of Scc2 to the centromere was independent of the tDNAs.

**Figure 4.**
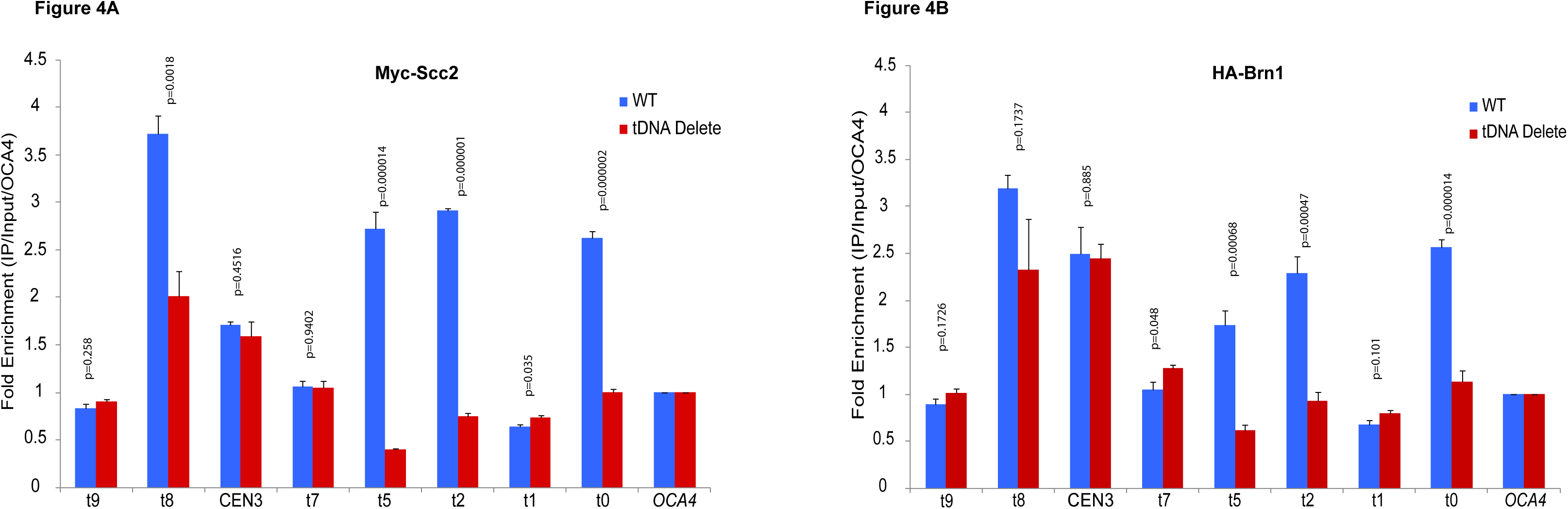
Scc2 and Brn1 binding at tDNAs on chromosome III. A) ChIP-qPCR mapping of Myc-Scc2. The data show the distribution of Scc2 at specific sites along chromosome III in the wild type and tDNA delete strain. The data are the results of two independent crosslinks from which four IP’s were performed. For each amplicon, the fold enrichment compared to input was first calculated and the data were then normalized to the *OCA4* locus. An unpaired t-test assuming unequal SD was used to test for significance of differences between the wild type and tDNA delete strain. B) ChIP-qPCR mapping of HA-Brn1, condensin. Fold enrichment and statistical significance was calculated in the same way as for the Scc2 ChIP and normalized to the *OCA4* locus.

Scc2, in association with Scc4 helps recruit the SMC proteins to chromatin (D’Ambrosio et al., 2008; Kogut et al., 2009). Condensins localize to tDNAs and are necessary for the clustering of tDNAs in the nucleus (D’Ambrosio et al., 2008; Haeusler et al., 2008). We therefore mapped the binding of condensins at tDNAs on chromosome III using the HA-tagged Brn1 subunit. In wild type cells, the Brn1 profile was very similar to that previously observed for Scc2 with significant binding of Brn1 at specific tDNAs. Correspondingly, the binding of the condensins was significantly reduced at these sites upon deletion of the tDNA promoters (Figure 4B).

### Chromosome mobility on the tDNA-less chromosome

TFIIIC binding sites and tDNAs are described as chromosome organizing clamps because of their consistent association with specific landmarks within the nucleus (Haeusler and Engelke, 2006). The localization of tDNAs with the kinetochore is dependent upon condensins while the interactions of tDNAs with nuclear pores are dependent upon cohesins. These associations likely help tether the chromosome. Since loss of tDNAs from chromosome III led to a decrease in SMC proteins from these sites we wondered if this loss would affect chromosome tethering and mobility of the chromosome. To assess mobility we fluorescently-labeled specific sites on chromosome III, and used these to monitor chromosome mobility in the wild type and the tDNA deletion chromosomes. The location of a point on the chromosome was mapped in three-dimensional space over a defined period of time in relation to another point within the nucleus-the spindle pole body (marked with the Spc29-RFP fusion protein)- and mobility was characterized by mean square distance analysis (MSD) (Dion et al., 2012; Mine-Hattab and Rothstein, 2012; Verdaasdonk et al., 2013). Six chromosomal loci across chromosome III were assayed (Figure 5). These loci were tagged by inserting LacO arrays at these sites and monitored using a LacI-GFP fusion protein mediated fluorescence. Time-lapse movies of individual unbudded cells in the G1 phase of the cell cycle were imaged over the course of 10 minutes. Using this information, MSD curves were generated for each locus in both the WT and tDNA delete strain (Supplementary Figure 4). For the wild type chromosome III, *CEN3* was the most constrained locus (Rc=415 nm), with loci located further from the centromere exhibiting greater mobility. For example, *LEU2*, which is approximately 30kb from the centromere, had an Rc of 522nm while *HMR*, which is approximately 180kb from the centromere, had an Rc value of 688nm. This is consistent with previous data showing that the location of a locus in relation to the centromere is critical in determining its mobility, with loci closer to the centromere displaying decreased mobility compared to loci farther from the centromere (Albert et al., 2013; Tjong et al., 2012; Verdaasdonk et al., 2013). Out of the six loci assayed, none of the loci showed a statistically significant change in mobility following the loss of tDNAs (p values range from 0.15 to 0.75). The data indicate that tDNAs are not major determinants in constraining chromosome segment motion or that they are a subset of factors involved and the redundancy precludes observation of their function.

**Figure 5.**
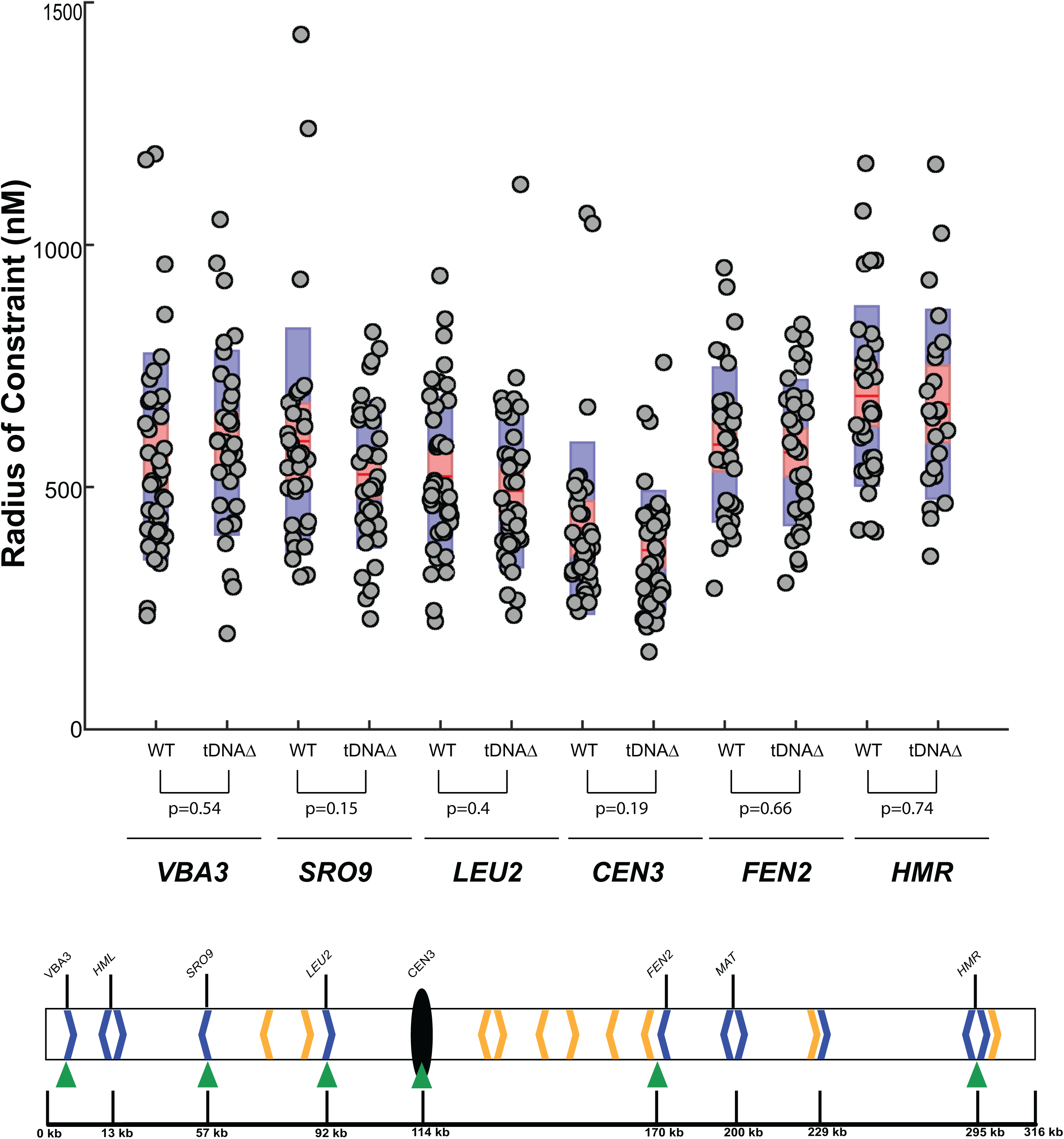
Effect of tDNA deletion on chromosome III mobility. Mean square displacement analysis of seven loci along chromosome III in wild type and tDNA delete strains are shown. Box plots represent the data obtained from the MSD experiments. Components of the boxplot are as follows; red line represents the mean, pink bar is the 95% confidence interval, purple bar is the standard deviation, and the grey dots represent individual values obtained from each cell analyzed. The green arrowheads beneath the chromosome III schematic show the locations of the loci assayed. The radius of constraint (Rc) measurement was calculated from MSD graphs that were generated over the course of a 10-minute time-lapse movie. These experiments are the result of time-lapse images taken from at least 35 cells per locus assayed. A t-test was used to determine significance of differences observed between the wild type and tDNA delete strain for each loci.

### tDNAs are not required for proper chromatin folding

Transfer RNA genes have been proposed to affect chromatin fiber folding via the clustering of dispersed tRNA genes. The promoters in tDNAs are the binding site for the transcription factor TFIIIC and foci comprised of multiple TFIIIC-bound sites have been proposed to function in chromatin looping and folding (Duan et al., 2010; Haeusler and Engelke, 2004, 2006; Iwasaki and Noma, 2012; Noma et al., 2006; Pombo et al., 1999; Raab et al., 2012; Thompson et al., 2003). If tDNAs are major drivers of chromatin folding and looping, then elimination of these loci from an entire chromosome should lead to changes in the folding of the chromatin fiber or result in changes in chromosome packaging in the nucleus. We set out to determine the detailed three-dimensional organization of chromosome III lacking functional tDNAs. We used a modified chromosome conformation capture technique called Micro-C XL (Hsieh et al., 2015; Hsieh et al., 2016). We chose Micro-C XL over HiC because it can capture both short length 3D interactions as well as long-length interactions and the method is not dependent upon the presence of restriction sites along the DNA. In brief, yeast cells were first cross-linked with formaldehyde and DSG, and chromatin was then fragmented into mono-nucleosomes via micrococcal nuclease digestion. Cross-linked, digested chromatin was ligated to capture chromosomal interactions. Size-selected ligation products were then purified and subjected to paired-end high-throughput sequencing. Sequencing reads were mapped back to the reference genome to determine the interacting regions of the chromosome, as previously described (Supplementary Figure 5). Overall, Micro-C maps for wild type and tDNA mutant strains both exhibited previously described features of yeast chromosome folding, with no difference in chromatin folding between the tDNA delete and wild type strains. For example, ∼2-10kb contact domains (CIDs/TADs) encompassing ∼1-5 genes were observed, across all 16 chromosomes, in the wild type strain. Inspection of the chromosome III in the tDNA delete cells showed that these domains persisted even upon loss of the tDNAs (Figure 6). There was no significant change in the contact frequency versus genomic distance in the two strains, indicating no local chromatin decondensation or change in chromatin looping interactions. Thus, tDNAs do not appear to be responsible for the general folding of the chromatin fiber.

**Figure 6.**
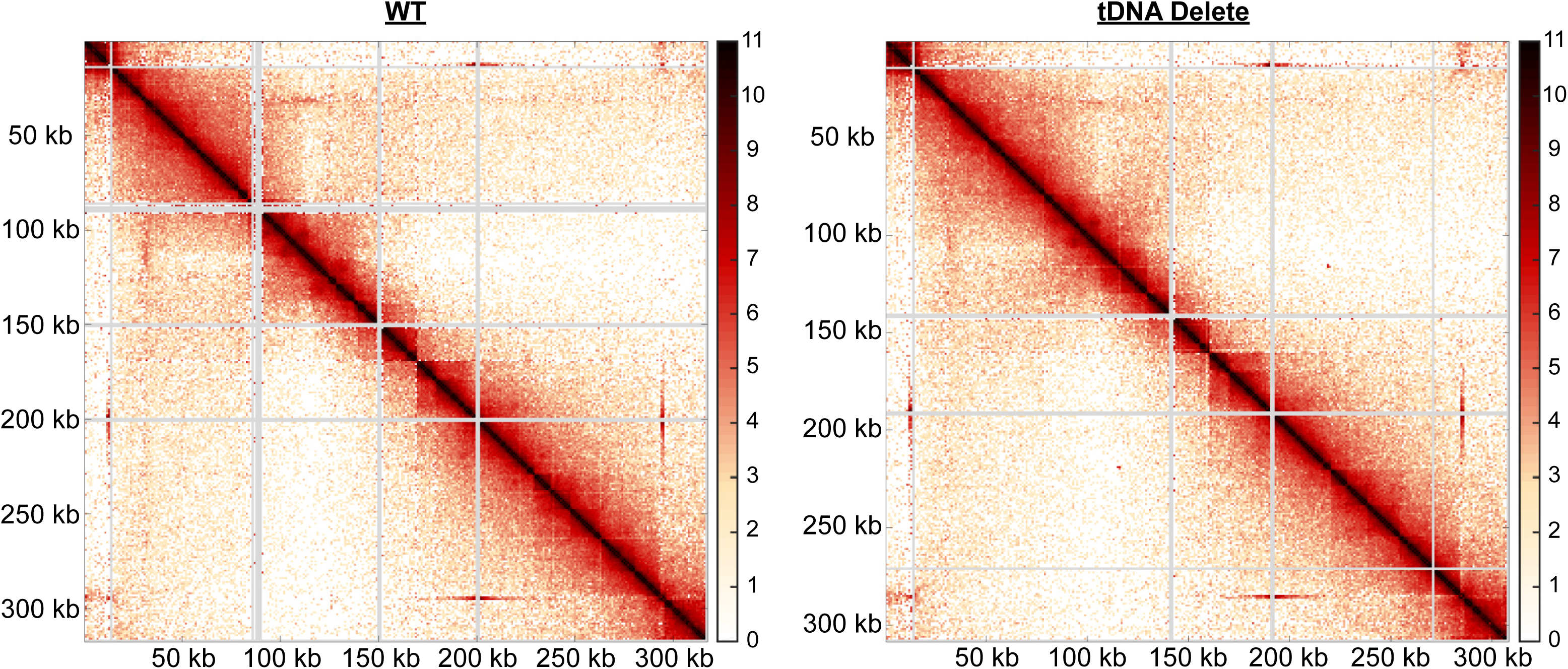
Micro-C Interaction plots of chromosome III. The heat maps show the interactomes for both wild type and tDNA delete strains for chromosome III. The X and Y-axes show the positions along the chromosome. Areas of red denote regions of high densities of internucleosomal interactions, while lighter colored areas denote decreased interaction.

### tDNAs affect CEN-CEN interaction frequency

While the overall folding of the chromatin fiber of chromosome III was not altered in the absence of tDNAs, and although Scc2 levels were unchanged at centromeres, Micro-C analysis identified changes in contact frequency at centromeres (Figure 7). The 16 centromeres in yeast are in close physical proximity to one another and cluster adjacent to the spindle pole body (Guacci et al., 1997; Jin et al., 1998; Jin et al., 2000). These CEN-CEN interactions are readily captured by 3C methods including HiC and Micro-C XL (Dekker et al., 2002; Duan et al., 2010; Hsieh et al., 2015), and are recapitulated in this study in the W-303 strain background. Interestingly compared to the wild type strain, the centromere of chromosome III in the tDNA delete strain showed an increased frequency of interactions with the other centromeres. Focusing on the 50kb pericentric region of each chromosome, we found that most CEN-CEN interactions were minimally affected by the loss of chromosome III tDNAs. For instance, interactions between the chromosome XVI centromere and the remaining centromeres showed that interactions between *CEN16* with the majority of centromeres remained unchanged, but that there was a ∼20% increase in interaction strength between *CEN16* and *CEN3* when chromosome III lacked tDNAs (Figure 7B).

**Figure 7.**
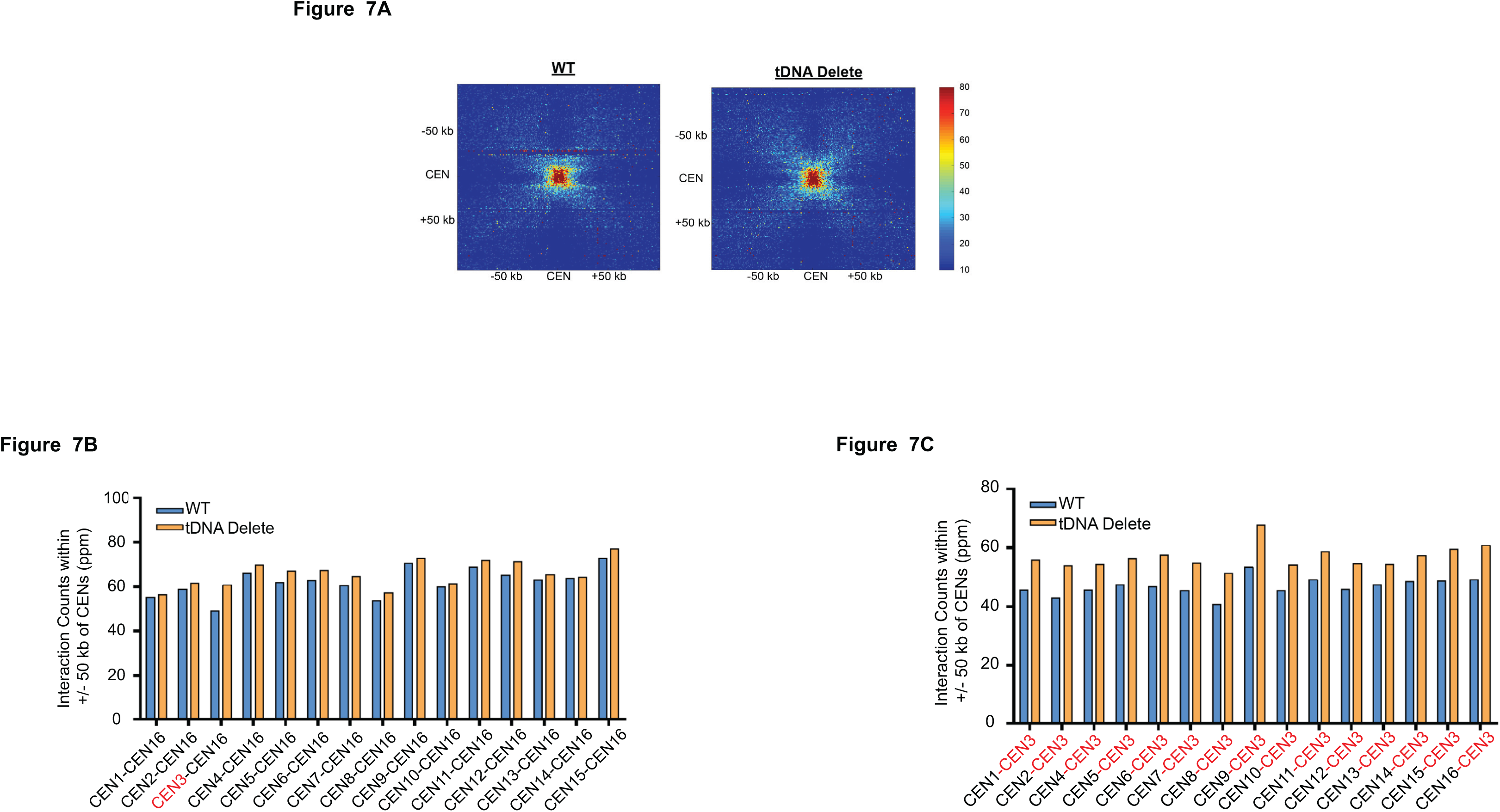
Micro-C analysis of the centromeres. A) Deletion of tDNAs on chromosome III leads to an increase in *CEN3* interaction with all other centromeres in the genome. Micro-C experiment analysis of *CEN-CEN* interactions is shown. The heat maps show a piled alignment of all centromeres. Interaction frequencies are denoted by the colored bar to the right of each heat map. B) The graphs are a quantification of *CEN-CEN* interactions. The graph examines the interaction of *CEN16* with all other centromeres. The x-axis is the interaction counts of a 50 kb segment centered on each centromere (in parts per million). C) The graph examines the interaction of *CEN3* with all other centromeres. The x-axis is the interaction counts of a 50 kb segment centered on each centromere (in parts per million). The increase in the CEN3-CEN interactions in the tDNA deletion strain was significantly higher (p=1.22×10^-14^) compared to values of all CEN16-CEN interactions (excluding CEN16-CEN3).

This increase in *CEN3* interaction was not confined to *CEN16*. When the same analysis was performed using *CEN3* as an anchor, we observed increased frequency of interactions between *CEN3* and all of the other chromosomal centromeres in the tDNA delete strain (Figure 7C). Most of the interaction counts increased approximately 20% compared to WT, with the highest increase seen at C*EN3-CEN9*. The increase in the CEN3-CEN interactions in the tDNA deletion strain was significantly higher (p=1.22×10^-14^) compared to values of all CEN16-CEN interactions (excluding CEN16-CEN3). These results show that upon deletion of all tDNAs across chromosome III, inter-chromosomal interactions increase between *CEN3* and the other centromeres, suggesting that functional tDNAs likely antagonize CEN-CEN associations during interphase.

### tDNAs play a role in *HML-HMR* long-range association

The silent loci *HML* and *HMR* reside on chromosome III separated by approximately 300kb along the linear chromosome. However, the *HML* locus, located 11kb from *TEL3L*, is in close three-dimensional proximity to the *HMR* locus, located 23kb from *TEL3R*. This long-range interaction has previously been detected using both live-cell microscopy and HiC analysis (Dekker et al., 2002; Kirkland and Kamakaka, 2013; Miele et al., 2009) and we recapitulate this finding in the Micro-C experiment with the wild type strain (Figure 8A). Comparing wild type cells to the tDNA delete strain, we noticed that the interaction of *HML* with *HMR* was slightly altered in the tDNA delete strain. In wild type cells, there was an interaction between *HML* and *HMR* and this interaction zone became less defined and more diffuse upon deletion of the tDNAs and a slightly increased interaction frequency was observed across a broader region of chromosome III. While *HMR* still interacted with *HML* in the deletion strain, it appeared to also display interactions with other loci (including *TEL3L)*. Similarly, the segment containing *HML/TEL3L* showed increased interactions with *TEL3R* rather than being restricted to interacting with sequences at *HMR*. These results suggest that deletion of chromosome III tDNAs subtly perturbed *HML-HMR* long-range interactions.

**Figure 8.**
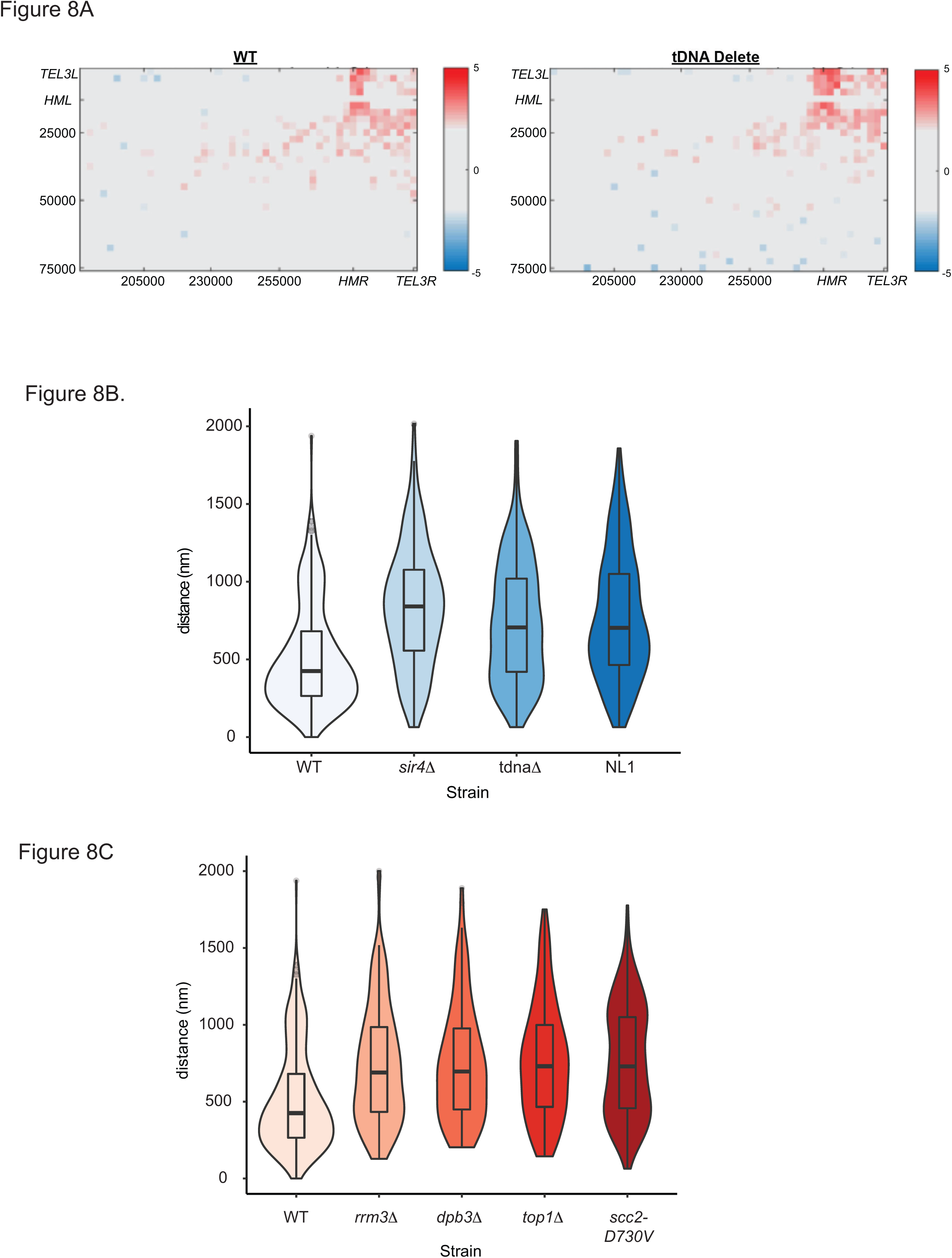
Long-Range *HML-HMR* association. Deletion of tDNAs on chromosome III leads to a change in *HML-HMR* interaction as measured by Micro-C. Heat maps display the interaction profile between segments on chromosome 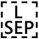 III that include *HML* and *HMR* (obtained from the Micro-C data). Increased interactions are denoted by red and decreased interactions are denoted by blue. The data are displayed in a log2 format. The x and y axes denote the area of the chromosome displayed on each axis of the heat map. B) Deletion of tDNA t0 leads to perturbation of *HML-HMR* long-range association. The violin plots show data of the distances between *HML*::TetR-YFP and *HMR*::CFP-LacI foci in asynchronously growing strains. Mann-Whitney U-Test were performed to determine statistical significance between wild type and the various mutants. Wild type (n=305), *sir4*Δ (n=134) (p=6.7 e^-16^), tDNA t0Δ (n=317) (p=3.1 e^-14^) or t0Δ:NL1 (n=330) (p=2.2 e-^16^) strains. The dark line in the middle represents the median distance. The data for *sir4Δ* are shown as a control and are the same as those in (Kirkland and Kamakaka, 2013). C) Replication–repair proteins are necessary for *HML-HMR* interactions: Violin plots of the distance between TetR-YFP and CFP-LacI foci in a given wild type or mutant strain are shown. rrm3Δ (n=208) (p=4.8 e^-12^), dpb3Δ (n=134) (p=1.7 e^-10^) top1Δ (n=139) (p=4.2 e^-12^) scc2D730V (n=188) (p=1.9 e^-14^). The data for the *scc2D730V* allele are simply shown as a control and are the same as those in (Kirkland and Kamakaka, 2013).

Given that Micro-C measures population averages of stable long-range interactions we decided to measure *HML-HMR* interactions in live cells using fluorescence microscopy. We wished to determine if the tDNA located adjacent to *HMR* influenced *HML-HMR* interactions. We generated a strain with multiple Lac operator sequences inserted adjacent to *HMR* (at the *GIT1* gene) and multiple copies of the Tet operator sequences inserted adjacent to *HML.* Expression of the fusion proteins CFP-LacI and YFP-TetR in this strain enabled us to visualize these loci in living yeast by fluorescence imaging. The distance between *HML* and *HMR* was then measured in wild type and a strain lacking the t0 tDNA (Figure 8B). We found that in wild type cells, *HML* was in close proximity to *HMR* and the median distance between these loci was 425nm. Consistent with our expectations, deletion of Sir proteins resulted in separation of these loci, with the median distance increasing to 840nm, validating this assay. Importantly, when we eliminated the t0 tDNA, this led to an increase in the distance between *HML* and *HMR* compared to wild type cells, with the median distance between *HML* and *HMR* shifting to 706nm upon deletion of the tDNA. Given the presence of outliers in the data we used a Mann-Whitney U-Test to determine statistical significance between wild type and the various mutants. With an n of approximately 300 cells for the wild type and tDNA delete strain we observed a p value of 3.1 e^-14^ showing that the differences observed were statistically significant. Closer analysis of the plot also indicates that upon deletion of the tDNA there is heterogeneity in the distances between the two loci and there is a continuum of values. Thus there are cells where the two loci are in very close proximity as well as cells where the two loci are very far apart and cells where the loci are at intermediate distances. This might help explain the fact that the difference observed by Micro-C is not large.

The tDNA t0 is necessary for the recruitment of cohesins to the silenced loci and the SMC proteins are necessary for long-range *HML-HMR* interactions (Kirkland and Kamakaka, 2013). However not all tDNAs are equivalent in their ability to recruit cohesins to the silenced loci (Donze and Kamakaka, 2001; Dubey and Gartenberg, 2007). We therefore inquired if tDNAs that are unable to recruit cohesins are able to restore long-range association between *HML* and *HMR*. We replaced the *HMR* tDNA^THR^ (t0) with tDNA^THR^ (NL1) from chromosome XIV. This tDNA has identical sequence to the t0 tDNA in the body of the gene and therefore has the identical BoxA and BoxB promoter sequence and spacing as tDNA t0. However sequences flanking this tDNA are distinct and the NL1 tDNA is unable to recruit/bind cohesins (Donze and Kamakaka, 2001; Dubey and Gartenberg, 2007). When we replaced a 300bp t0 tDNA-containing fragment with a 300 bp NL1 tDNA-containing fragment we found that the NL1 tDNA was not able to robustly restore long-range *HML-HMR* interactions, suggesting that tDNA mediated cohesin loading might be necessary for these long-range interactions.

### Replication fork pausing at tDNA mediates long-range chromosomal interactions

We had previously shown that double strand repair proteins help deposit cohesins to the silenced loci, which then led to homology dependent long-range interactions between *HML* and *HMR* (Kirkland et al., 2015; Kirkland and Kamakaka, 2013). Since we had discovered that a specific tDNA played a role in the clustering of *HML* and *HMR*, we wished to know the mechanism by which this phenomenon occurred. Replication fork pausing/stalling is observed at many tDNAs. This results in the deposition of γ-H2A at the tDNA and this is necessary for fork recovery from the pause/stall (Achar and Foiani, 2017; Azvolinsky et al., 2009; D’Ambrosio et al., 2008; Deshpande and Newlon, 1996; Kirkland and Kamakaka, 2013; Lengronne et al., 2004; Szilard et al., 2010). Rrm3 and topoisomerases play a role in the recovery of stalled replication forks at protein bound sites in the genome such as tDNAs (Achar and Foiani, 2017; Azvolinsky et al., 2006; Ivessa et al., 2003). Therefore we analyzed the effect of these mutants on *HML-HMR* long-range association. The data show that deletion of Rrm3 as well as mutants in the DNA polymerase-ε subunit Dpb3 and the topoisomerase Top1 lead to a statistically significant decrease in *HML-HMR* long-range association (Figure 5B and figure legend). Thus the presence of a tDNA as well as normal Rrm3, Top1 and Dpb3 function are necessary for the establishment or maintenance of the long-range association between *HML* and *HMR*.

### tDNA presence enhances epigenetic gene silencing at clustered *HML-HMR*

Since the t0 tDNA located adjacent to the silenced *HMR* domain is necessary for the long-range clustering of the silenced domain, we wondered if reduction in clustering had any effect on gene silencing. We asked whether tDNA mediated loss of *HML-HMR* interactions affected gene silencing at *HML* and *HMR*. Silencing can be assayed by insertion of reporter genes within or immediately adjacent to the silenced domains. In wild type yeast when a reporter gene is inserted immediately adjacent to these loci, the genes is metastably silenced. A cassette containing an H2B (*HTB1*) promoter driving *HTB1-EYFP* was integrated to the right of *HML* while a cassette containing the *HTB1* promoter driving *HTB1-ECFP* was integrated to the left of *HMR*. In addition, on chromosome XV, a cassette containing an *HTB1*promoter driving *HTB1*-mCherry was integrated as a control euchromatic marker (Mano et al., 2013). The *HTB1*-mCherry gene is active in all cells in the population. The *HML::YFP* and the *HMR::CFP* reporter genes are present immediately outside of *HML* and *HMR* but reside in a region bound by Sir proteins (Oki and Kamakaka, 2005; Ruben et al., 2011). These genes adopt one of two expression states, either active or silent. For visualization, single cells were placed on microfluidic plates and monitored continuously by fluorescence microscopy. Fluorescent signal from each individual cell was recorded every 40 minutes over a period of ∼24 hours. This allowed us to trace the lineage of each daughter from the founder cell and score the cells according to the expression of the reporter genes at *HML* and *HMR*. Cell lineage trees were traced and each cell in the lineage was assigned a positive or negative value for expressing each reporter as it underwent cell division (Figure 9A).

**Figure 9.**
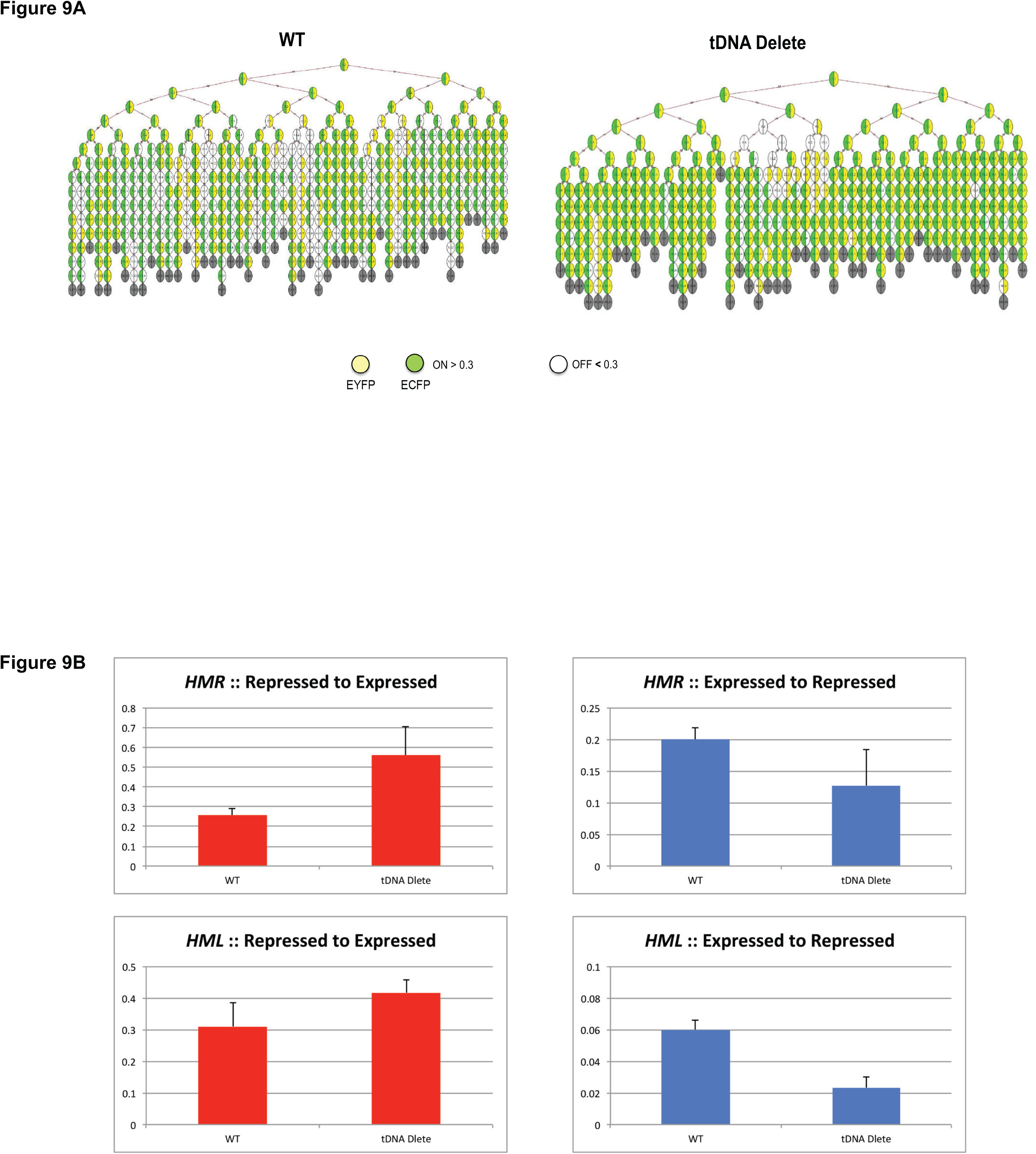
Silencing of reporter genes at *HML* and *HMR*. tDNAs on chromosome III modulate silencing of reporter genes at *HML and HMR*. Representative lineage trees of the different strains that were analyzed are shown. Wild type refers to a strain containing all tDNAs on chromosome III. tDNA delete refers to a strain lacking any tDNAs on chromosome III. The expression of *HML::EYFP* or *HMR::ECFP* in each generation of cells was monitored, quantitated and is indicated by the presence of their respective colors in the cells of the tree. B) Deletion of tDNAs on chromosome III leads to a change in the maintenance of silencing at *HML* and *HMR*. The graphs quantify the changes in expression state of *HML::EYFP* and *HMR::ECFP* between generations in the different genotypes studied. Expressed to repressed transitions identify reporters that were expressed in one generation but not expressed in the next. Repressed to expressed transitions represent reporter genes that were not expressed in one generation but expressed in the next.

We initially analyzed the silencing of the reporter genes in the wild type strain. Consistent with previous data (Mano et al., 2013), reporters at *HML* and *HMR* were regulated such that the reporters maintained their activity state over many generations and occasionally switched to the opposite expression state. Once they switched they maintained the new state for several generations. Furthermore, when one reporter was active the other was also more likely to be active suggesting long-range coordination between *HML* and *HMR* though this coordination is not absolute.

We next investigated silencing of the reporters in a strain where chromosome III lacked all the tDNAs. In this strain, the reporter at *HMR* was active more often compared to the wild type strain. While the effect was not as pronounced, the same effect was also observed at *HML* (Figure 9A tDNA delete panel). Furthermore, the silenced state was less stable, and switched to the active state more often. This suggests that silencing at these loci is influenced by the tDNAs.

While expression states at both *HML* and *HMR* were stably inherited, the transcriptional state did flip in daughter cells (Figure 9B). An expressed to repressed transition was a less frequent event compared to the repressed to expressed transitions regardless of genotype. This is not entirely surprising since the reporter genes were inserted immediately outside of the two silencers in a zone where the silent state is metastable (Valenzuela et al., 2008; Valenzuela et al., 2006). However, when analyzing the repressed to expressed transitions, we saw a discernible difference in the frequency of the expression of the reporter genes at *HMR*. The full tDNA delete strain showed an increased frequency of cells undergoing the transitions at *HMR* compared to wild type cells and the inverse was seen for the expressed to repressed transition (Suppl. Table 3).

Taken together, the data suggest that deletion of tDNAs on chromosome III had an effect on the ability of *HMR* to interact with *HML* and diminution of this clustering led to an alteration in the stability of the silenced state at these loci.

## Discussion

tDNAs are middle repetitive DNA sequences scattered across all 16 chromosomes and their primary function is the synthesis of tRNAs. In this manuscript, we show that tDNAs affect local chromatin structure, which then impinges on chromosome architecture. tDNAs 1) affect chromatin structure by maintaining local nucleosome free regions along the fiber and precisely positioned nucleosomes immediately outside of the tDNAs, 2) recruit cohesins and condensins 3) affect nuclear architecture by influencing centromere clustering and 4) alter heterochromatin clustering leading to changes in the fidelity of inheritance of gene silencing.

The binding of specific proteins such as CTCF to a site on the DNA can affect nucleosome positions over long distances (Fu et al., 2008). Nucleosome depletion at the gene and positioned nucleosomes flanking the gene is a hallmark of tDNAs (Cole et al., 2012; Dhillon et al., 2009; Dion et al., 2007; Harismendy et al., 2003; Moqtaderi and Struhl, 2004; Nagarajavel et al., 2013; Oki and Kamakaka, 2005; Roberts et al., 2003; Yuan et al., 2005). Our data show that loss of the tDNA promoters’ only affect nucleosome positions in the immediate vicinity of the tDNA. The nucleosome positioning effects mediated by the tDNA bound transcription factors TFIIIC and TFIIIB are not transmitted over long distances.

### tDNAs, SMC proteins and chromatin folding

The SMC proteins are involved in higher order chromosome organization in all eukaryotes and have been extensively mapped. tDNAs are binding sites for all three classes of SMC proteins (cohesin, condensin and repairsin), the SMC protein loaders Scc2 and Scc4 and the meiotic Rec8 SMC protein (Blat and Kleckner, 1999; D’Ambrosio et al., 2008; Glynn et al., 2004; Klein et al., 1999; Kogut et al., 2009; Laloraya et al., 2000; Lindroos et al., 2006; Noma, 2017). Given these intimate connections between tDNAs and the SMC proteins our data indicate that loss of the tDNA promoters does lead to loss of SMC proteins from tDNAs but this effect is tDNA specific since we do not see a loss of SMC proteins from centromeres or Scc2 from other sites in the genome. Surprisingly the loss of Scc2 and Brn1 from tDNAs does not affect chromatin folding. While clustering of tDNAs in the nucleus (as measured by fluorescence microscopy) is dependent upon the SMC proteins (Haeusler and Engelke, 2006; Noma et al., 2006) the precise contribution of tDNAs in this process remained unclear. Our Micro-C analysis of chromosome III suggests that tDNAs are unlikely to be the key drivers of chromatin looping and folding or tethering to nuclear substructures since we did not observe any change in contact frequency across the entire chromosome. Furthermore, we did not observe any discernible effect on chromosome loss rates or chromosome mobility. It is likely that tDNA independent SMC protein binding sites masks the tDNA-mediated effects. SMC proteins bind only half of the tDNAs in the nucleus and only a third of the SMC protein binding sites localize at or near tDNAs (D’Ambrosio et al., 2008). The lack of phenotype would also be consistent with previous data that showed that a reduction in the levels of the SMC proteins does not affect the properties of the chromosome arm (Heidinger-Pauli et al., 2010). Recently a synthetic yeast chromosome III was generated and characterized (Annaluru et al., 2014; Mercy et al., 2017). The synthetic chromosome lacks repetitive sequences such as TY elements, LTRs and tRNA genes. The 3D structure of this chromosome was determined using HiC and the data show that there were no major differences between this chromosome and the wild type chromosome except for a shortening of the length. While this chromosome lacks multiple elements the three-dimensional folding data are consistent with our conclusions from the Micro-C analysis of the same chromosome lacking only tDNAs. While it is possible that redundancy of structural elements masks tDNA-mediated effects on chromatin folding it is also possible that chromatin folding is driven by underlying DNA sequence and not tDNA mediated interactions. The yeast chromosomes have isochores with G-C rich, gene rich R-band segments alternating with AT-rich G-band segments (Dujon, 1996; Sharp and Lloyd, 1993), which exhibit different functional properties (Blat et al., 2002; Dekker, 2007). Chromosome III has a G-C segment from 20 to 100 kb on the left arm followed by an A-T rich central segment from 100 to 200kb on the right arm and then a second G-C rich segment from 200 to 290kb on the same arm. In this scenario, the underlying A-T rich DNA sequence likely plays a dominant role in the three dimensional folding of chromatin. tDNAs are often syntenic along chromosomes (Raab et al., 2012; Raab and Kamakaka, 2010) and it is possible that these positions have been selected for optimal gene activity rather than being involved in chromatin loop formation (Gehlen et al., 2012). Thus while the A-T rich isochore is structurally and functionally distinct (Baudat and Nicolas, 1997; Dekker et al., 2002; Gerton et al., 2000) and is the region rich in tDNAs (See Figure1) our results would suggest that the tDNAs do not play a significant role in either tethering of this isochore or the overall folding of this segment. The tDNA clustering observed by microscopy could simply be a function of linear proximity of tDNAs along the chromatin fiber.

### tDNAs and centromere clustering

Chromosome tethering to nuclear substructures enables nuclear organization (Gehlen et al., 2012; Taddei et al., 2010) and centromeres and the telomeres along with their associated proteins play a key role in this process (Andrulis et al., 2002; Bupp et al., 2007; Bystricky et al., 2004; Duan et al., 2010; Gartenberg et al., 2004; Guacci et al., 1997; Jin et al., 2000; Mekhail et al., 2008; Schober et al., 2009; Taddei and Gasser, 2004; Tjong et al., 2012). All sixteen centromeres cluster together in a ring around the membrane-embedded spindle pole body. The centromeres are tethered to the spindle pole body via direct interactions between kinetochore-associated proteins and the spindle pole body associated microtubules in interphase (Bystricky et al., 2004; Dekker et al., 2002; Guacci et al., 1997; Jin et al., 2000). Other factors are likely to influence this phenomenon but remain unknown. tDNA density is almost 2 fold higher in the pericentric region of *S. cerevisiae* chromosomes including chromosome III (Snider et al., 2014) (see Figure 1). While tDNAs have been shown to help tether centromeres to the spindle axis during mitosis (Snider et al., 2014), in interphase nuclei, the loss of tDNAs results in increased interactions between the clustered centromeres. The physical presence of tDNAs in the pericentric region could interfere with the close packaging of centromeres during interphase. This could be due to transcription-mediated effects since tRNA genes are highly active. In *S. pombe*, mutations that reduce tDNA transcription result in increased tDNA association with the centromere and increase chromosome condensation during mitosis. Furthermore, tDNA association with centromeres increases when the genes become inactive (Iwasaki et al., 2010). Thus, tDNA clustering at transcriptionally active RNA pol III transcription factories near the centromeres could hinder closer centromere-centromere interactions during interphase while a decrease in tDNA transcription during mitosis could help tether centromeres to the spindle axis during mitosis (Snider et al., 2014).

An alternative though not mutually exclusive possibility is based on the observation that transcriptionally active tDNAs interact with nuclear pores (Casolari et al., 2004; Chen and Gartenberg, 2014; Ruben et al., 2011). It is thus possible that there is a competition between pericentric tDNA-nuclear pore interactions in opposition to centromere-centromere interactions. In this scenario, the loss of tDNA tethering to the nuclear pore would enable the centromere greater freedom of movement thus enabling closer centromere-centromere interactions.

### tDNA effects on *HML-HMR* interactions and the inheritance of gene silencing

Gene silencing is primarily a function of the Sir proteins though numerous other factors influence the process (Gartenberg and Smith, 2016). Proto-silencers are sequence elements that on their own are unable to silence a gene, but when located near a silencer increase the efficiency of silencing (Fourel et al., 2002; Lebrun et al., 2001). Our demonstration that the tDNA affects silencing of a reporter adjacent to the silent *HMR* domain suggest that tDNAs function as proto-silencers. Our data suggest that tDNA mediated clustering of silent loci might be important in the silencing of these loci and the loss of long-range association might reduce the efficient inheritance of the silent state. This is analogous to the observations that gene clustering at active chromatin hubs and transcription factories increases the efficiency of transcription as well as the data showing that telomere clustering increases the efficiency of silencing at sub-telomeric sequences (Gasser et al., 2004).

This unexpected observation also raises the question of how might tDNAs influence long-range *HML-HMR* interactions. tDNAs, including the tDNA next to *HMR*, are sites of replication slowing/pausing (Admire et al., 2006; Azvolinsky et al., 2009; Deshpande and Newlon, 1996; Ivessa et al., 2003; Lemoine et al., 2005; Wang et al., 2001). The tDNA adjacent to *HMR* is a site of replication fork pausing (Kitada et al., 2011; Szilard et al., 2010). We recently showed that long-range *HML-HMR* interactions require homologous sequences to be present at these loci (Kirkland et al., 2015; Kirkland and Kamakaka, 2013) and we now show that mutations in replication coupled homologous recombination repair proteins including the SMC proteins, Rrm3, Top1 and Dpb3 lead to a reduction in *HML-HMR* interactions. Based on the accumulated data we would posit that replication fork slowing/pausing results in the deposition of γH2A and SMC proteins at tDNAs followed by a homology search leading to *HML-HMR* interactions. The re-formation of silenced chromatin following replication precludes the eviction of γH2A (Keogh et al., 2006) thereby stabilizing SMC protein binding, which then maintains the long-range *HML-HMR* association. The tDNAs thus help initiate a network of interactions mediated by the SMC proteins and the Sir proteins leading to *HML-HMR* association and chromosome folding. We would like to posit that a series of transient interactions during replication aid in the setting up of the final optimal nuclear architecture found in the interphase nucleus.

In conclusion, tDNAs primarily affect local chromatin structure. Each tDNA affects nucleosome positions and protein binding in its immediate vicinity. These local perturbations functionally and structurally interact with neighboring regulatory regions resulting in tDNA mediated pleiotropic effects. In some instances tDNAs affect the expression of neighboring pol II transcribed genes by the phenomenon of local tgm silencing. In another context tDNA mediated replication pausing result in the establishment of long-range heterochromatin interactions, which then influence the inheritance of silencing states at these loci.

## Acknowledgements

This work was supported in part by a grant from the NIH to RTK (GM078068) and (T32-GM008646) to JK, OH and KW. This study was funded in part by the Intramural Research Program of the National Institutes of Health (NICHD). We thank the NHLBI Core Facility for paired-end sequencing of the micrococcal nuclease libraries. The Functional Genomics Laboratory, UC Berkeley sequenced the ChIP-Seq libraries on an Illumina HiSeq4000 at the Vincent J Coates Laboratory at UC Berkeley, supported by an NIH S10 OD018174 Instrumentation Grant.

## Materials and Methods

### Yeast strains and primers

Table S1 and S2 list the yeast strains and the primer sequences that were used in this study.

### MNase-Seq

MNase-Seq experiments were carried out as previously described (Cole et al., 2012). In brief, isolated nuclei were digested with MNase to mono-nucleosomes. Paired-end sequencing libraries were prepared (Illumina). Paired reads (50 nt) were mapped to the reference genome (SacCer2) using Bowtie-2 (Cole et al., 2014; Langmead and Salzberg, 2012; Ocampo et al., 2016). For analysis of nucleosome occupancy (coverage) at tDNAs, both across the genome and on chromosome III, tDNAs were aligned on their start sites or at the deletion points. Data sets were normalized to their genomic average, set at 1, using only DNA fragments in the 120 to 180 bp range. In one experiment, mono-nucleosomal DNA was gel-purified, but not in the replicate, in which short fragments (< 120 bp) derived from digestion of the TFIIIB-TFIIIC complex at tDNAs (Nagarajavel 2013) were observed. The MNase-seq data are available at the GEO database:(GSE98304 https://www.ncbi.nlm.nih.gov/geo/query/acc.cgi?token=wnynwaoqvnktfmb&acc=GSE98304)

### ChIP-Seq and RNA-Seq

Chromatin immunoprecipitation reactions were performed essentially as described above but elution of the precipitated DNA from Protein A/G beads was carried out with two successive washes in 175ul of 0.1M NaHCO_3_/1% SDS. 50ul of each input sample was diluted to 350ul with the elution buffer. NaCl was added to a final concentration of 0.2M and cross-links were reversed with an overnight incubation at 65C in a Thermomixer. All samples were treated with 60ug of RNAase A (Sigma) at 37C for 60’ followed by a Proteinase K (Roche) treatment at 50C for 60’. DNA was purified with a successive phenol chloroform and chloroform extraction followed by precipitation with 2 volumes of ethanol and 50ug of glycogen (Roche).

The ChIP and Input DNA was spun, washed with 70% ethanol and re-suspended in deionized water. DNA quantitation was performed using a Qubit dsDNA HS Assay kit prior to confirmation by qPCR.

Libraries for ChIP-Seq were prepared at the Functional Genomics Laboratory, UC Berkeley and sequenced on an Illumina HiSeq4000 at the Vincent J Coates Laboratory at UC Berkeley.

For RNA-Seq, yeast strains JRY2334 and JKY690 were grown in duplicate in 50ml YPD to a cell density of 6-7×10^6^ cells/ml, spun, washed in 25ml PBS, divided into 4 aliquots per culture and transferred to 1.5ml microfuge tubes. Cell pellets were flash frozen in liquid N2 and transferred to −70C. RNA, library preparation and sequencing for RNA-Seq were performed by ACGT Inc. Wheeling, IL.

Transcript abundances were estimated using Kallisto (Bray et al., 2016). Differential analysis of gene expression data was performed using the R package Sleuth (Pimentel et al., 2017). Likelihood ratio test and Wald test were used to identify the differentially expressed genes (false discovery rate adjusted p-value (or q-value) < 0.05 in both tests). Since the likelihood ratio test does not produce any metric equivalent to the fold change, we used the Wald test to generates the beta statistic, which approximates to the log2 fold change in expression between the two conditions.

Sequence data have ben deposited in the GEO database.

https://www.ncbi.nlm.nih.gov/geo/query/acc.cgi?token=krihsykczdmbpox&acc=GSE106_250

### ChIP

ChIP-qPCR experiments on all Brn1 and Scc2/4 were performed as previously described (Dhillon et al., 2009; Kirkland and Kamakaka, 2013). In brief, yeast cells of a strain of interest were inoculated and grown overnight in 300 ml of YPD media to an OD of 1-2. These cells were then fixed in 1% formaldehyde for a duration of 2 hours at room temperature. The reaction was then quenched with glycine, and the cells were spun down and washed in 1X PBS. The cross linked cells were then flash frozen in dry ice and stored at −70^0^C. In preparation for IP, the cells were thawed on ice, broken apart by bead beating, and sonicated to achieve a desired chromatin size of ∼300 bp. Once the size of the chromatin was checked, cell debris was cleared from the sample by high-speed centrifugation. The cross linked, sized chromatin was split into 2 samples and IP’s were done overnight in the presence of both an antibody to the protein of interest as well as pre-blocked A/G-Sepharose beads at 4°C. 50 µl of input chromatin was also taken from each IP sample prior to addition of the antibody. Chromatin elution was done using 10% Chelex 100 (Bio-Rad) along with proteinase K treatment. After elution, both input and IP DNA were quantitated via a Picogreen fluorescent quantification assay (Invitrogen). For each qPCR reaction, input DNA was run in triplicate and IP DNA was run in duplicate. An equal amount of input and IP DNA was used in each individual reaction. The enrichment for a given probe was then calculated as IP/Input, and was further normalized to the OCA4 locus. The results of each ChIP-qPCR are comprised from two independent crosslinks per strain assayed, and for each crosslink two independent IPs were done.

### Mean Squared Distance Analysis

Mean-squared distance analysis was carried out as previously described (Hediger et al., 2002; Heun et al., 2001; Verdaasdonk et al., 2013). In brief, we built strains that contained a 64x lacO array at specific points along chromosome III. We then integrated a cassette containing an spc29-RFP fusion protein elsewhere in the genome. This protein is an essential kinetochore protein, and therefore serves as a marker for the spindle pole body. The spindle pole body served as a fixed point to which we could measure the movement of our GFP tagged loci in 3D space over a period of 10 minutes. Z-stack images of the cells were taken every 30 seconds during the time-lapse, and the data are used to calculate the radius of constraint using the equation: <(X_t_ – X_t+Δt_)2>. MSD curves were generated for each locus in both the WT and tDNA delete strain (Supplementary Figure 2). The plateau of the MSD curve was used to calculate the radius of constraint (Rc) for each locus. This analysis was performed in no less than 35 cells per genotype assayed. The data were plotted in “NotBoxPlots” (source code obtained from https://github.com/raacampbell/notBoxPlot)

### HML-HMR Colocalization analysis

Distance assays between *HML* and *HMR* was performed as previously described (Kirkland and Kamakaka, 2013). Fluorescence microscopy was performed on live yeast cells after growing the cells in YMD with Leucine, uracil, tryptophan, lysine, adenine and histidine. Cells were grown to an Od A600 of approximately 0.6. Cells were washed in YMD, placed on YMD-agar patches on slides, and imaged. Microscopy was performed using an Olympus xi70 inverted wide-field microscope with DeltaVision precision stage using a Coolsnap HQ2 camera and a 100x/1.4 oil objective. The 20 image stacks for each image were acquired with a step size of 200nm using the appropriate wavelength for CFP, YFP, GFP or mCherry. The acquisition software used was softWoRx3.7.1. The images were cropped using Adobe Photoshop. For the distance analysis between *HML* and *HMR*, the distance between the yellow and cyan dots were calculated in nanometers using the “measure” tool in three dimensions. The measured distances were loaded into R software (www.r-project.org) and the data were plotted as a box plot. The box includes the middle 50% of the data with the line in the box being the median value. The data presented are the sum of at least two independent strains.

### Single Cell Expression Analysis

Single cell expression analysis was performed as previously described (Mano et al., 2013). Briefly, cells were grown in YPD at 30C and placed in a microfluidics device. Time-lapse photos of growing cells were recorded using an Axio Observer Z1 microscope using a 40x objective. The ECYP and EYFP fluorescence intensities were normalized to the highest level of fluorescence observed and the Euchromatic mCherry signal.

### Micro-C

Micro-C was performed as previously described (Hsieh et al., 2015). The detailed method have been described (Hsieh et al., 2016). In brief, this technique provides nucleosome level resolution of all of the interactions occurring across the genome by using MNase digestion in lieu of a restriction enzyme as in traditional Hi-C techniques. The interactome data were deposited in the GEO database GSE98543 (https://www.ncbi.nlm.nih.gov/geo/query/acc.cgi?acc=GSE98543)

### Antibodies

Antibodies used in ChIP were as follows; Scc2-Myc: anti-myc 9E10 (Abcam) = 5 μl, Brn1-HA: anti-HA HA.11 (Covance) = 5 μl.

**Supplementary Figure 1) Nucleosome occupancy plots at five tDNA deletion points on chromosome III.** MNase-seq data for wild type and tDNA delete were normalized to the genomic average (= 1). Coverage plots are shown using all DNA fragments in the 120 to 180 bp range. The reference point (0) is the nucleotide marking the 5’-end of the deletion on chromosome III. Upstream of the deletion point at 0, the DNA sequence is the same in wild type and the tDNA delete chromosome III. Downstream of the deletion point, the DNA sequences are different. The black arrow shows the location and orientation of the tDNA in wild type chromosome III. Meaningful plots cannot be made for two tDNAs (tP(AGG)C and tS(CGA)C), because they were moved to another chromosome. Two other tDNAs (tM(CAU)C and tK(CUU)C) are present in S288C strains but are naturally absent in W-303 strains, including the strains used here. The wild type profile is in orange and the tDNA delete profile is in blue.

### Figure Legends

**Supplementary Figure 2) Deletion of tDNAs does not lead to general changes in chromatin structure at RNA pol II transcribed genes on chromosome**

**III.** Comparison of global nucleosome phasing on chromosome III in wild type (blue line) and tDNA delete (red line) cells: average nucleosome dyad positions on 106 RNA Pol II-transcribed genes on chromosome III. These genes cover most of chromosome III. The genes were aligned on their transcription start sites (TSS set at 0). The average nucleosome dyad density is set at 1.

**Supplementary Figure 3**) **ChIP-Seq graphs showing the distribution of Scc2 at specific sites along chromosome III.** tDNAs are marked by arrowheads. Graph 1 and 2 are independent ChIP-Seq data from the wild type strain. Graphs 3 and 4 are independent ChIP-Seq data from the tDNA delete strain.

**Supplementary Figure 4) Summary graphs for MSD analysis for each genotype**

The graphs summarize the results of MSD time-lapse experiments for the 8 loci assayed. Both WT (blue) and tDNA 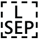 delete (red) are shown on each graph. Time-lapse images were taken over a 10 minute period, z-stack images were taken every 30 seconds in 200 nm increments. The distance between the GFP marked locus 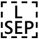 of interest and spc29-RFP spindle pole body protein was calculated at each 30-second interval. Mean distance traveled for each time-point was used to create the MSD curves. At least 35 cells were assayed for each genotype.

**Supplementary Figure 5) Whole genome interaction plots for Micro-C experiments that were done in this study.** The heat maps show full genome interactomes for both wild type and tDNA delete strains. On the x-axis, the chromosomes are displayed from chr I to chr XVI going from left to right. On the y-axis, the chromosomes are displayed from chr I to chr XVI going from top to bottom. Areas of red denote increased interaction, while lighter colored areas denote decreased interaction.

**Supplementary Table1:** Strain list with genotypes

**Supplementary Table2:** Sequences of PCR primers used in this study

**Supplementary Table3:** Statistical analysis of differences in expression of *HML::EYFP* and *HMR::ECFP* in the wild type and tDNA delete strains.

